# Plant-soil nitrogen, carbon and phosphorus content after the addition of biochar, bacterial inoculums and N fertilizer

**DOI:** 10.1101/2020.02.18.954941

**Authors:** Irina Mikajlo, Bertrand Pourrut, Brice Louvel, Jaroslav Hynšt, Jaroslav Záhora

## Abstract

The use of biochar in combination with mineral or biological amendments in order to improve its influence on soil-plant properties has received growing attention. The changes of N, C and P content in *Lactuca sativa var. capitata* aboveground plant biomass and soil after the addition of beech wood biochar combined with the addition of bacterial inoculums (Bacofil and Novarefm) and N fertilizer have been studied using spectrophotometry methods. Pots were filled with the arable soil from the plots in protection zone of water sources (Březová nad Svitavou, South Moravia, Czech Republic). Biochar with inoculums decreased plant growth in the first yield of Novaferm treatment and in both yields of Bactofil treatment. Increased plant biomass growth was observed with Novaferm addition in the second yield. Total N increase has been obtained in the plant aboveground biomass and soil of the treatments amended with inoculums and nitrogen fertilizer. The decrease of P content has been observed in plant aboveground biomass in the biochar amended samples.

## Introduction

Agricultural lands have been exposed to anthropogenically soil degradation resulted into its productivity loss, frequently caused by overfertilization and poor water management involving soil erosion [1]. Worldwide approximately 45% of arable soils are degraded [2] and annually 0.3–0.8 % of global arable land is considered improper for agricultural production [3]. In Czech Republic soil erosion caused by water, soil compaction and reduction in soil organic matter (SOM) are the most important types of soil degradation that were induced by past intensive farm practices [4].

Thus, soil restoration strategies have to be implemented aiming to mitigate soil degradation trends [5]. In the last decade, biochar (BCH) has attracted an increasing interested to improve the arable soil due to its potential agronomic benefits and carbon sequestration ability. BCH can have an effect on soil-forming processes that in turn determine the transformation, translocation and accumulation of soil constituents, changing soil pedogenic activity, morphology, and productivity from the long term perspective [6]. BCH could also play a role as soil conditioner for improving soil fertility and crop productivity [7,8]. Overall BCH influence on soil fertility depends on the amounts of carbon and macro or micronutrients derived from the BCH feedstock and pyrolysis temperature [9,10]. Though, improvements of global soil nutrient availability may take a certain period of time to be observed [9].

Soil amendment with a biochar has direct influence on soil microbial communities and their activity [11,12]. According to Martínez-Viveros et al. [13] plant growth-promoting bacteria (PGPB) create significant component of the soil microflora improving plant health along with productivity, while BCH with its porous structure of specific appropriate dimensions may serve as an effective inoculum carrier providing a protected habitat for bacteria, inaccessible for predators [14]. In addition, BCH can retain organic and inorganic nutrients from the feedstock and adsorb nutrients from root exudates thus successfully supporting inoculum development after its introducing into the soil [15].

In the other hand, BCH could reduce plant growth [16]. This phenomenom could be explained by the presence of toxic compounds due to charring process, like polycyclic aromatic hydrocarbons (PAHs), cresols, xylenols, formaldehyde, acrolein and volatile compounds such as pyroligneous acids (PA) [17] or the immobilization of nutrients by BCH [18,19], and especially nitrogen [11,20,21].

Some researchers have shown the interest of applying BCH in combination to mineral or organic amendments in order to mitigate potential BCH negative effects. Plant growth responses are significant when charcoal and fertilizers are combined, assuming a synergistic relationship [22,23]. According to Steiner et al. [23], BCH combined with N fertilizer is efficient for crop yield enhancement while reducing the N application. Considering the influence of BCH on soil microorganisms, the co-application of BCH with biological amendments could be also of interest. Several studies demonstrate the effect of inoculated BCH by bacterial inoculums mixed with fertilizers effect [8,24–26], however there are still missing data about their combined acting after the plant influence and persistence in soil. Moreover, there is also insufficient amount of studies regarding comparison of freshly applied BCH with the persisted in soil BCH after the second yield as the vast majority of research is based on one growing cycle investigation. In addition, the BCH influence is broadly dependent on the BCH feedstock and production temperature.

In this study, we investigated the influence of BCH combined with other soil amendments (bacterial inoculums and N fertilizers) on lettuce (*Lactuca sativa L.*) growth and soil-plant nutrient content (C, N and P) during two growing cycles.

## Materials and methods

### Soil sampling and preparation

Soil has been collected randomly from a permanent experimental Banín plots situated in the protection zone of underground drinking water source “Brezova nad Svitavou” (Czech Republic; 49°40.409’N, 16°27.545’E.) Soil was sampled with a spade according to Czech Technical Standard ISO 10 381-6 from the topsoil horizon (till 0-30 cm) in summer 2015. The soil is characterized as sandy loam type of luvisol **(Table 1)**. Fresh soil samples have been delivered to the laboratory where have been air-dried, homogenized and sieved through a 10-mm sieve.

**Table 1.**
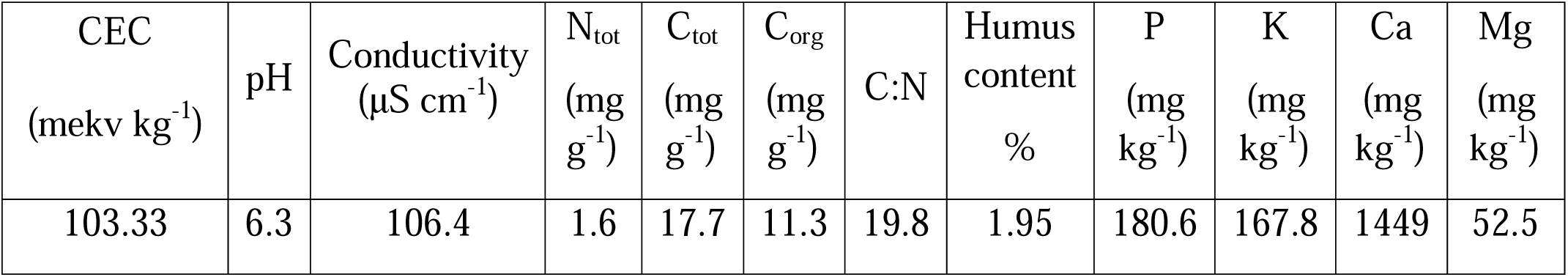

| CEC<br>(mekv kg<sup>-1</sup>) | pH | Conductivity<br>(μS cm<sup>-1</sup>) | N<sub>tot</sub><br>(mg g<sup>-1</sup>) | C<sub>tot</sub><br>(mg g<sup>-1</sup>) | C<sub>org</sub><br>(mg g<sup>-1</sup>) | C:N | Humus<br>content<br>% | P<br>(mg kg<sup>-1</sup>) | K<br>(mg kg<sup>-1</sup>) | Ca<br>(mg kg<sup>-1</sup>) | Mg<br>(mg kg<sup>-1</sup>) |
| --- | --- | --- | --- | --- | --- | --- | --- | --- | --- | --- | --- |
| 103.33 | 6.3 | 106.4 | 1.6 | 17.7 | 11.3 | 19.8 | 1.95 | 180.6 | 167.8 | 1449 | 52.5 |

According to Pokorný et al. [27] this loamy soil is characterised by lightly acidic pH, lower C_org_ content, low N_tot_ (0.1-0.3%), weak humus content, high P content, low Mg content which are typical for Czech soils. The conductivity index states on the increased salt content (60-120 μS cm^-1^).

### Biochar material

Beech wood biochar (*Fagus silvatica L.*) has been originated from Czech Republic (company BIOUHEL.CZ s.r.o.). It is characterized by the production at slow pyrolysis with the use of low temperature 470°C **(Table 2)**.

**Table 2.**
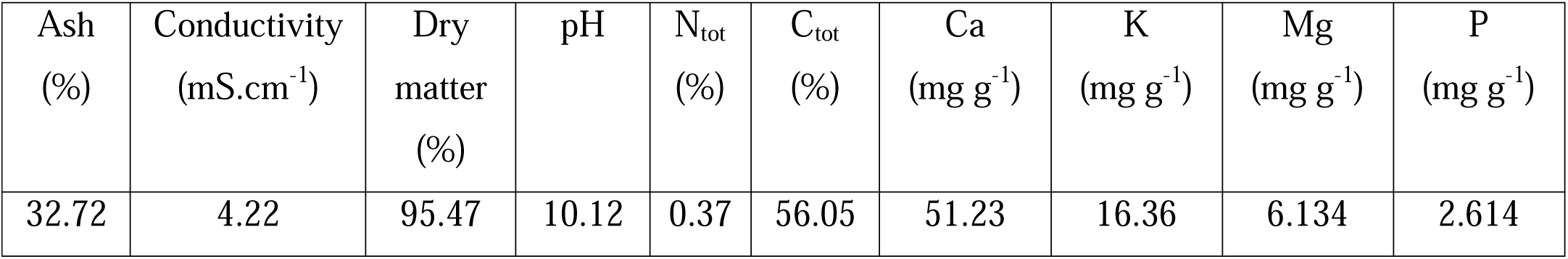

The C content may be characterized as the one obtained from the grass-derived biomass, rather than the wood-derived one (56.05%) with the high ash content, low P content and average C/N ratio of 151.5 [28]. It also exhibits an alkaline pH (10.12).

### Experimental design

Five different types of treatments including a control have been prepared **(Table 3)** that included four replications of each treatment resulting into twenty plastic square containers (10×10×11 cm) filled with 800 g of topsoil.

**Table 3.**
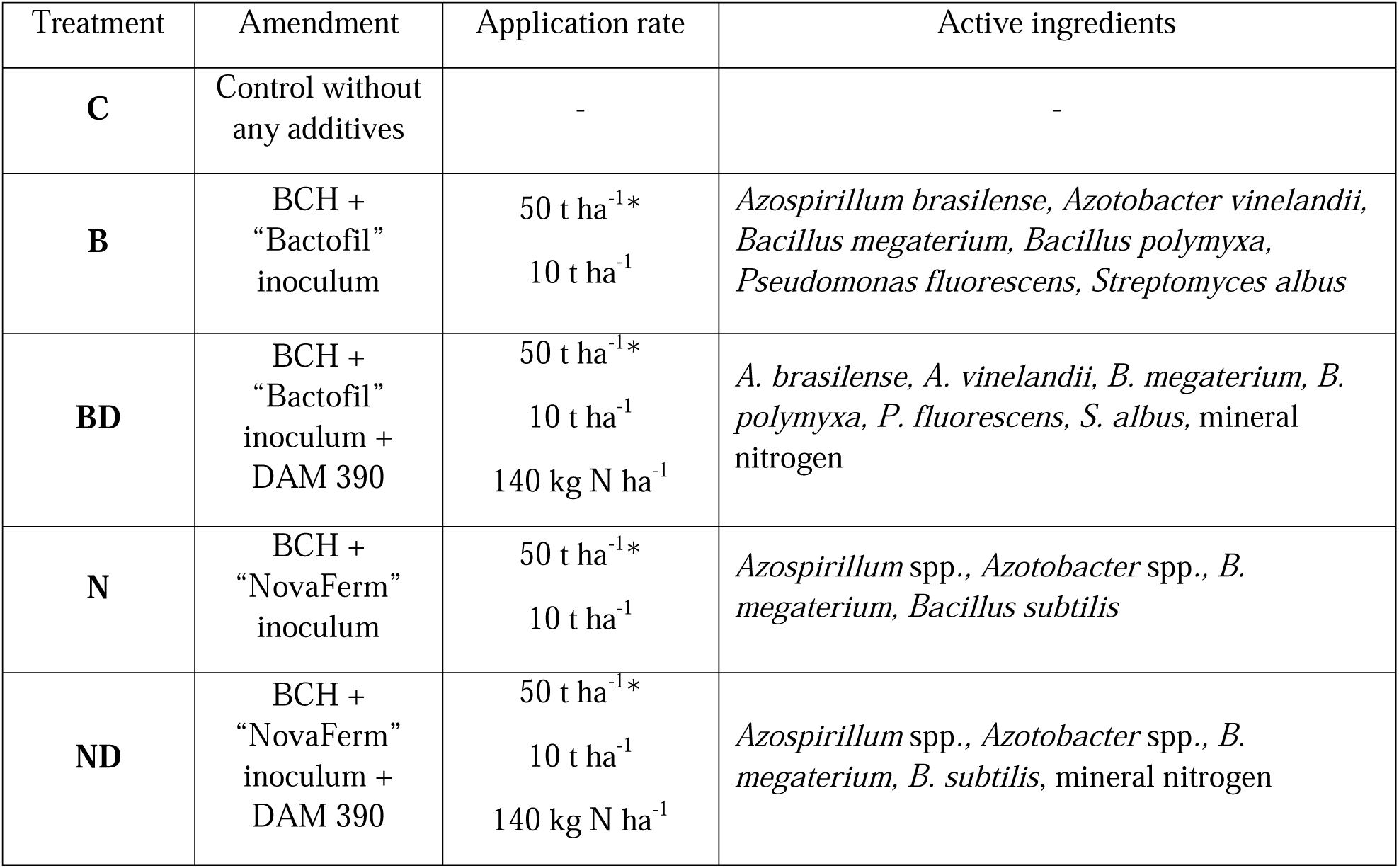

BCH was freshly applied in the quantity of 6.5% per pot with the first plant growing cycle. The BCH quantity used for the experiment was chosen as a high concentration in order to obtain distinguish results [22]. In the following experiment with second plant growing cycle, the same soil (e.g. with the same BCH) was reused again.

Pots were split into two groups. Half of them were inoculated with the commercial bacterial inoculums “Bactofil” (B) from BioFil Ltd (Budapest, Hungary) while the other half were inoculated with “NovaFerm” (N) from Nova Scienta Kft (Soltvadkert, Hungary) at the beginning of the experiment on lettuce (BBCH-scale: 13). Later (BBCH-scale 15-18), DAM 390 fertilizer has been added to half of the inoculated pots (BD and ND) at the dose recommended by the supplier (140 kg N ha^-1^). DAM 390 is a liquid fertilizer of ammonium nitrate with urea and with amide (N-NH_2_) nitrogen and ammonium (NH_4_-N) nitrogen suitable for fertilizing. It contains 30 % of nitrogen; the ratio of ammonium, nitrate, and amidic nitrogen is 1:1:2.

### Plant cultivation

*Lactuca sativa var. capitata L. cv. Kennedy* has been chosen as an experimental plant. Each pot contained one plant. The pots were randomly put in CLF PlantClimatics® growth chamber that was set to maintain temperatures of 22 °C by day and 19 °C at night, 65% humidity, with a day length of 12 h and light intensity of 380 μmol m^-2^ s^-*1*^ using automatic vents and supplementary heating. All the pots have been watered by adding 50 ml of deionized water every 2 days.

After one growing cycle, leaf biomass was harvested, weighted and prepared for nutrient analysis. Root biomass from the first growing cycle of plants was removed and experimental soils were homogenized again and re-filled into containers. The same lettuce plant has been seeded. The inoculation by “Bactofil” and “NovaFerm” additives and the application of DAM 390 fertilizer were done at the same BBCH-scale as in the case of the first plant growing cycle.

At the end of the experiment, soil and plant samples were collected and analysed for SOC, total N and P contents.

### Nutrient determination in soil and Lactuca sativa aboveground biomass

#### Plant and soil samples preparation

Lettuce aboveground biomass was cut and moved into plastic trays then dried at 40 °C to invariable weight. After the determination of total dry plant biomass, plants were grounded into fine powder utilising a knife mill (GM200, Retsch). Soil samples were cleared from plant residues, sieved through 2-mm sieve and transferred into plastic trays to dry at 40 °C in the oven.

#### Nutrient content analysis

500 mg of soil sample and 50 mg of dry plant sample were used for nutrient content determination.

Soil organic carbon (Corg) levels were determined with a spectrocolorimetric dosage according to a method of sulphochromic oxidation (NF X 31-109, 1993). In a hood, 5 mL of potassium dichromate (K_2_Cr_2_O_7_, 80 g L^-1^), then 7.5 mL of sulfuric acid (H_2_SO_4_, 95%) were added to a glass tube containing 500 mg of soil sample sieved to 250 µm, or 50 mg of dry plant sample crashed with a mill. Then, glass tube was stirred thoroughly and placed in a heating block (DK Heating Digester (Velp Scientifica®) at 140°C for 30 min. Following solution was cooled and was poured in a 100 mL graduated flask, where level was adjusted with osmosed water. With the next step after stirring, an aliquot of 20 mL was taken and centrifuged for 10 min (4500 rpm; 2000 g). 200 µl of all samples were placed in a microplate and the absorbance was measured at the wavelength λ= 585 nm using a spectrophotometer Multiskan® GO. Corg was expressed in g kg^-1^ released in the sample and was obtained after calculating the blank absorbance with referring to the calibration curve. Standard curve was made with potassium dichromate (K_2_Cr_2_O_7_, 80 g L^-1^) solution adding 1 g of glucose. Replicates were used for quality assurance, white and certified materials BCR® 129, IRMM, Belgium) were included in the analyses.

N and P were analysed in the same soil and plant extracts in accordance with the Kjeldahl digestion procedure, modified by Saha et al. [29] for nitrogen and in accordance to the ascorbic/molybdate method described in the French standard NFX 31-161 and in Joret and Hébert [30] for phosphorus. For N and P estimation in soil, 0.5 g of dry soil sample was weighted to a glass digestion tube with addition of 3.5 g of a catalyst composed by the mixture K_2_SO_4_, CuSO_4_ x 5 H_2_O, and TiO_2_ (proportion of 33:1:1). Then, samples were vortexed. Samples of plant aboveground biomass of 50 mg were weighted to a glass digestion tubes with adding 3.5 g of the same catalysts. A volume of 10 mL of concentrated (96.8%) H_2_SO_4_ and 10 mL of 30% H_2_O_2_ was successively added slowly to each tube and then mixed by swirling. Thereafter, the digestion tubes were placed on a digestion block (DK Heating Digester, Velp Scientifica), heated at 200 °C for 20 min and at 390 °C for 45 min. The samples were digested until the solution was clear green. Digests were cooled for 15 min at room temperature and diluted by adding distilled water to constitute a solution of 100 mL volume and measured by UV/VIS spectrophotometer (Multiskan® GO) at a wavelength λ = 660 nm.

In general, total Kjeldahl nitrogen appears to be the sum of free-ammonia and organic nitrogen compounds that are transformed to ammonium sulphate (NH_4_)_2_SO_4_, within the described digestion conditions. By the use of the certified reference sample (IRMM, BCR-129, Belgium) the exactness of the N and P analytical determinations were verified.

For the total P assessment, 1 ml of digest product was transported in a glass tube and a colouring reagent was added to the final solution (ammonium heptamolybdate tetrahydrate in H_2_SO_4_). Standard curve was prepared from orthophosphate, solutions, after they were measured spectrophotometrically (UV - 1800 Shimadzu) at wavelength λ = 825 nm. The total P has been expressed in mg g^-1^ released from samples and was obtained after calculating the blank absorbance and referring it to the calibration curve.

### Data analysis

Nutrients concentrations in soil and aboveground biomass of lettuce are expressed and presented as the means and standard deviations of replicates for each treatment within two plant growing cycles. Analysis of variance (ANOVA) was accomplished to estimate within- and between-two plant growing cycle differences concerning elements concentration in soil and including the aboveground biomass weight across the treatments. The normal distribution of data (Shapiro-Wilk test) and equality of variances (Bartlett test) were checked. When both tests proved conformity, Fisher statistics was considered for significance (*p* ≤ 0.05) and the Tukey (HSD) test was used for pair-wise comparisons of statistical groups. The Kruskal-Wallis test was performed for data that were not distributed normally. All statistic tests were conducted in XLSTAT software (AddinsoftTM software 2016).

## Results

### Lettuce growth/Plant biomass measurements

In the first growing cycle (G1) results ranged from 0.5 till 2.3 g DM **(Figure 1)**. No statistically significant differences have been found between the treatments and control (1.6 g DM). However, treatments amended with DAM fertilizer had significantly higher values (respectively 1.2 times in BD treatment and 1.4 times in ND treatment compared to the control).

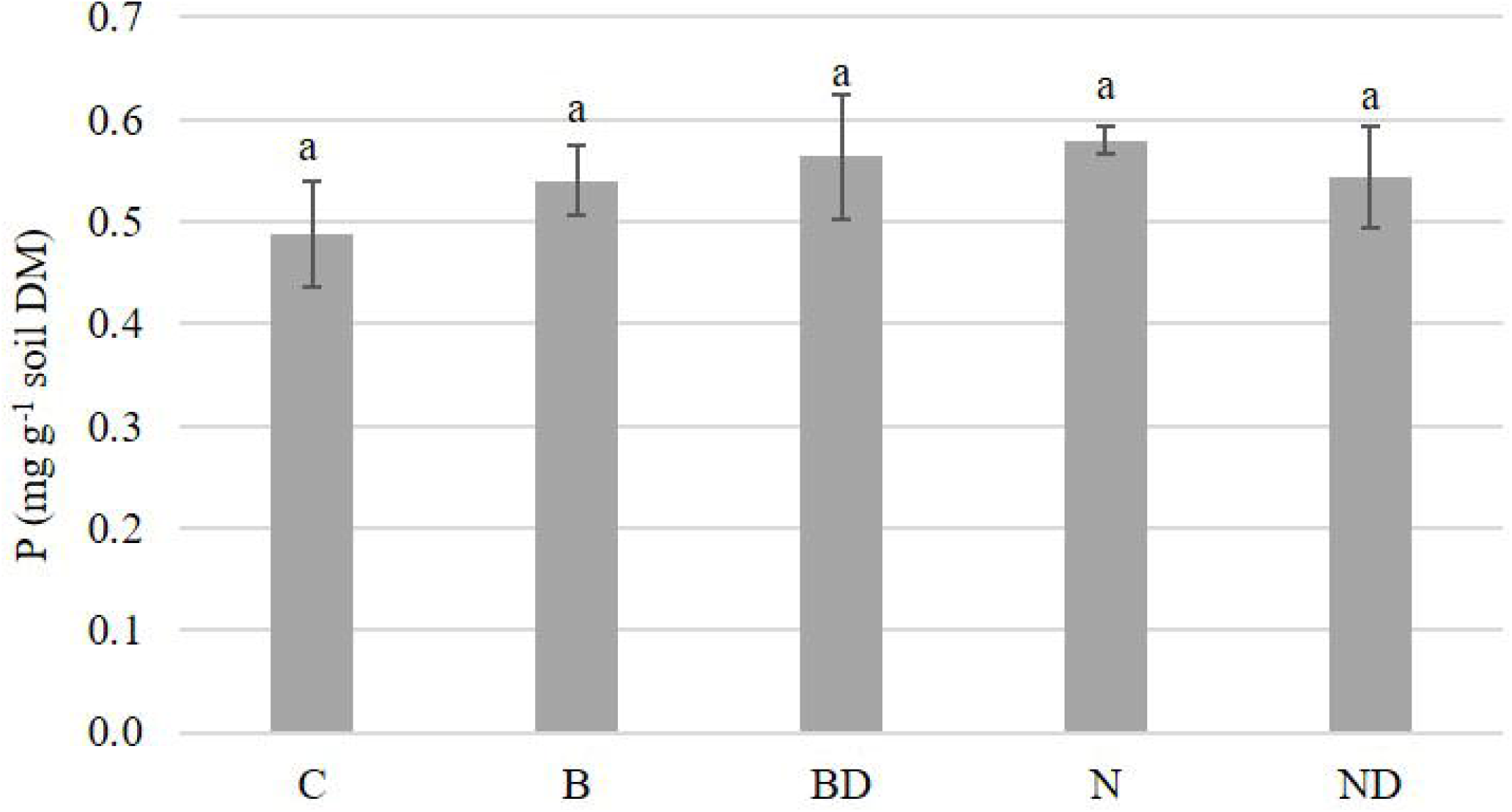

In the second growing cycle (G2) values have ranged from 0.5 till 4.1 g DM. No significant differences have been found between the C and B treatment, but the significantly increased values in BD treatment (by 6 times), ND treatment (by 7.2 times) and N treatment (by 8.3 times) respectively to the control plant.

Taking into comparison two growing cycles G1 and G2, no significant differences have been found within the treatments, except the Novaferm inoculated treatments, where N treatment showed significant increase by 8.2 times in G2 comparing to G1 and ND treatment by 1.6 times respectively.

### Total N in biomass/soil

In G1, N content values in lettuce biomass fluctuated from 12.3 mg g^-1^ leaves DM to 25 mg g^-1^ leaves DM **(Figure 2)**. No significant differences have been observed between the C, B, N and ND treatments, though BD value was significantly higher (2 times) from the other treatments except N treatment value. In G2, no significant differences have been detected within B, BD comparing to control soil (10.8 mg g^-1^ leaves DM), where N treatment differed from C, B and ND treatment, but not the BD treatment. ND value was significantly higher that all the other treatments and by 2.9 times comparing to the C.

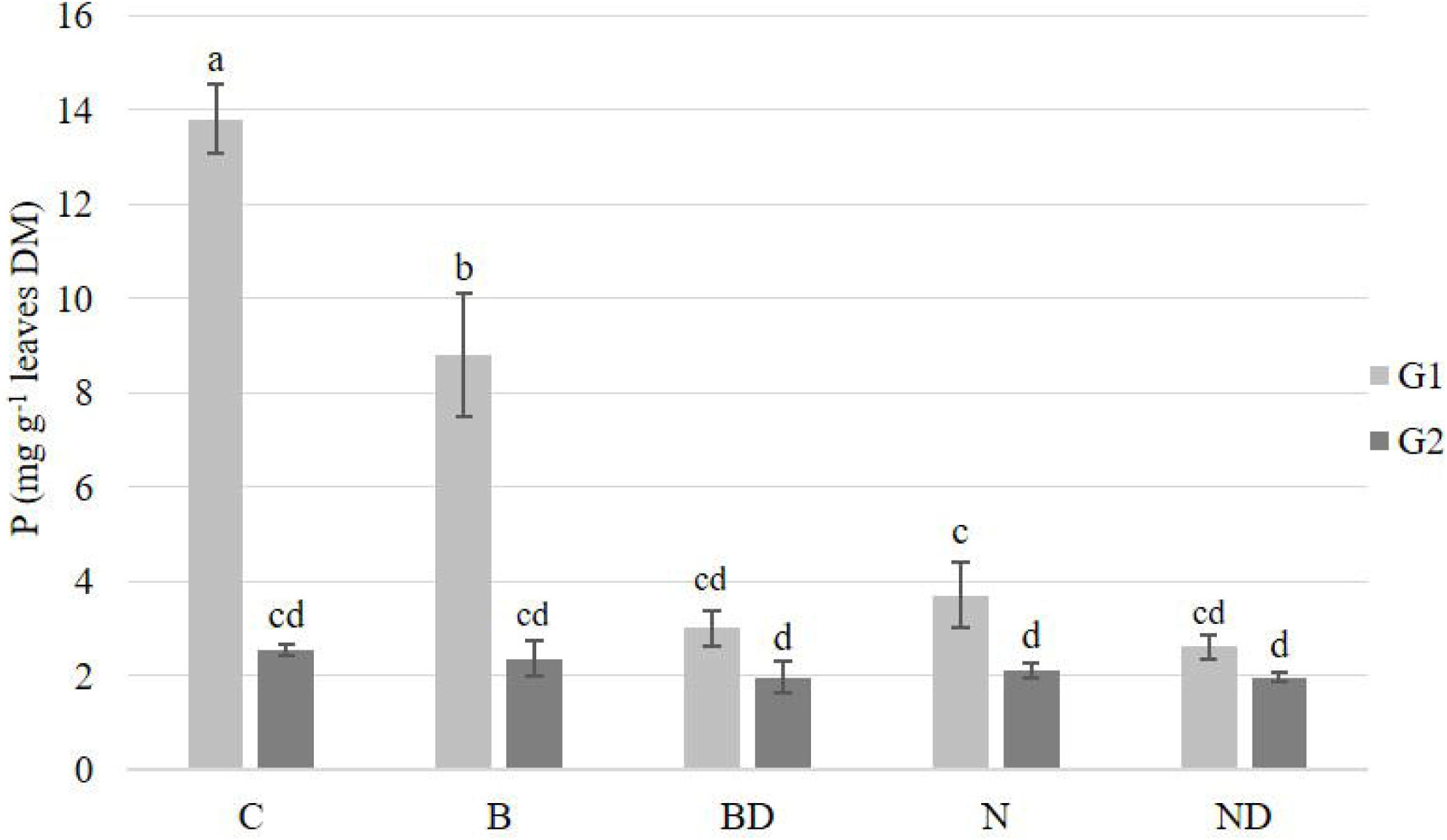

Comparing G1 and G2 reveals that no significant differences were observed that characterize the C, B and N treatments, although DAM amended treatments showed a decrease up to 34% in the case of its combination with Bactofil inoculum (BD) and an increase in 51.8% with the Novaferm inoculum combination (ND).

**Figure 3** shows that BCH amended treatments exhibited significantly higher N values in BD, N and ND treatments comparing to control soil (0.6 mg g^-1^ soil DM), though B treatment did not show any significant differences comparing to the rest of the treatments.

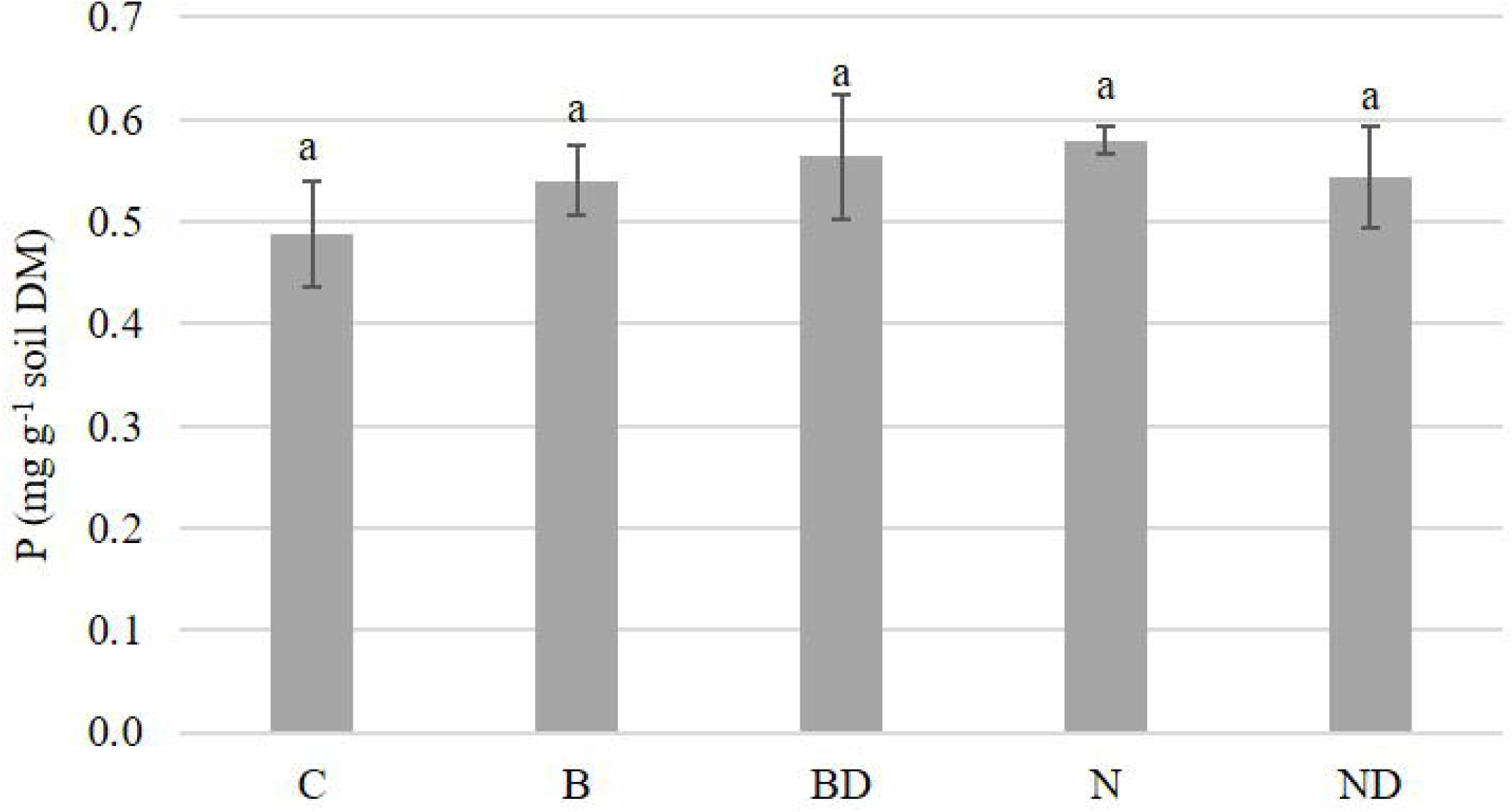

### Corg content in biomass/soil

No statistically significant differences have been detected in G1 and G2 regarding Corg content in the lettuce biomass **(Figure 4)**. Corg content fluctuated in G1 from 337-364 mg g^-1^ leaves DM and in G2 from 317-347 mg g^-1^ leaves DM.

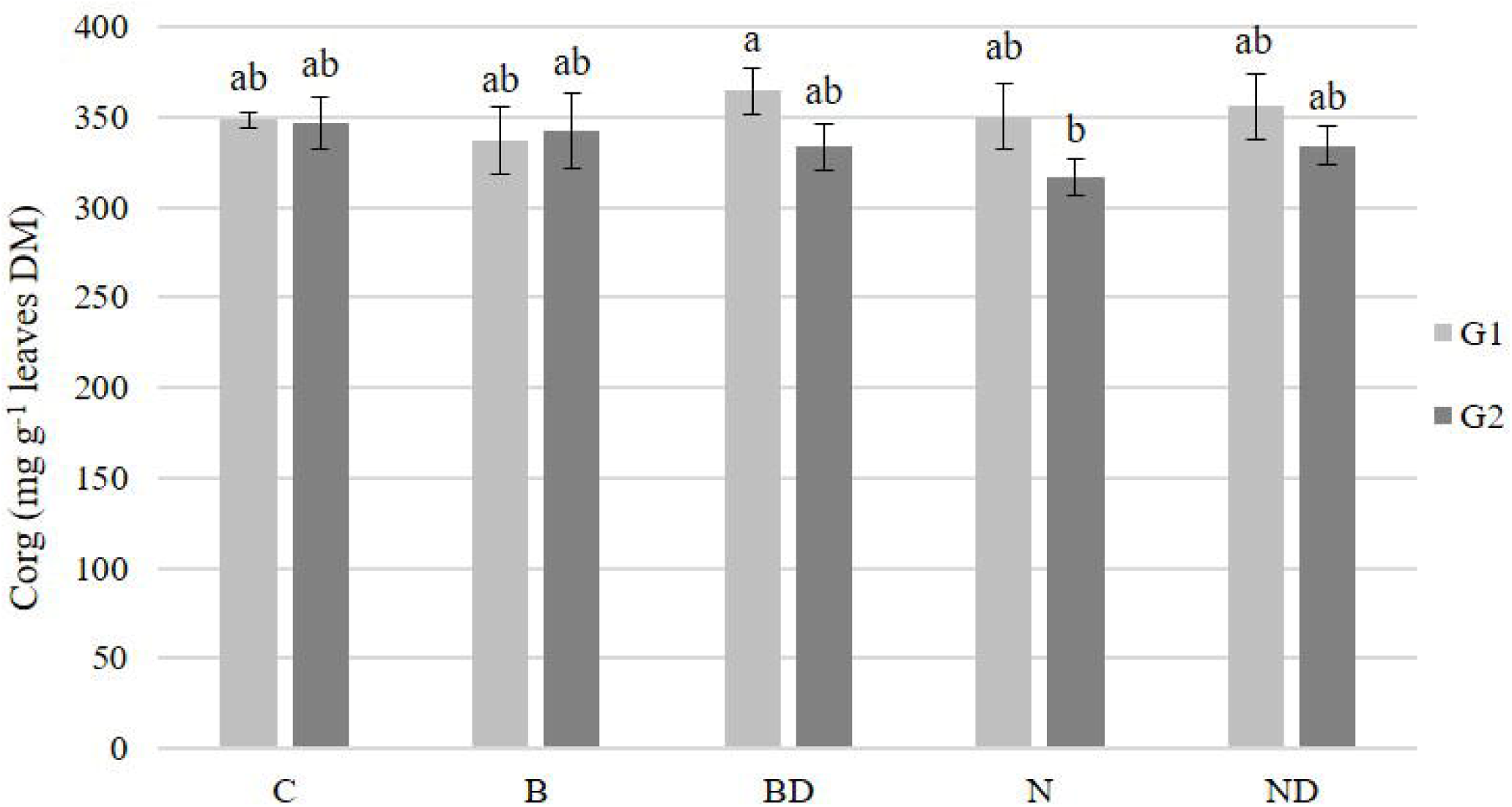

Soil Corg content has been detected in significantly higher amounts in the BCH amended treatments comparing to the control soil (8.7 mg g^-1^ soil DM).

No differences have been found between B, N and ND treatments itself, while BD treatment was 5 times higher comparing to control soil, but meantime having no significant difference with the N treatment **(Figure 5)**. B and ND treatment were significantly higher from control soil (3.7 and 3.6 times respectively).

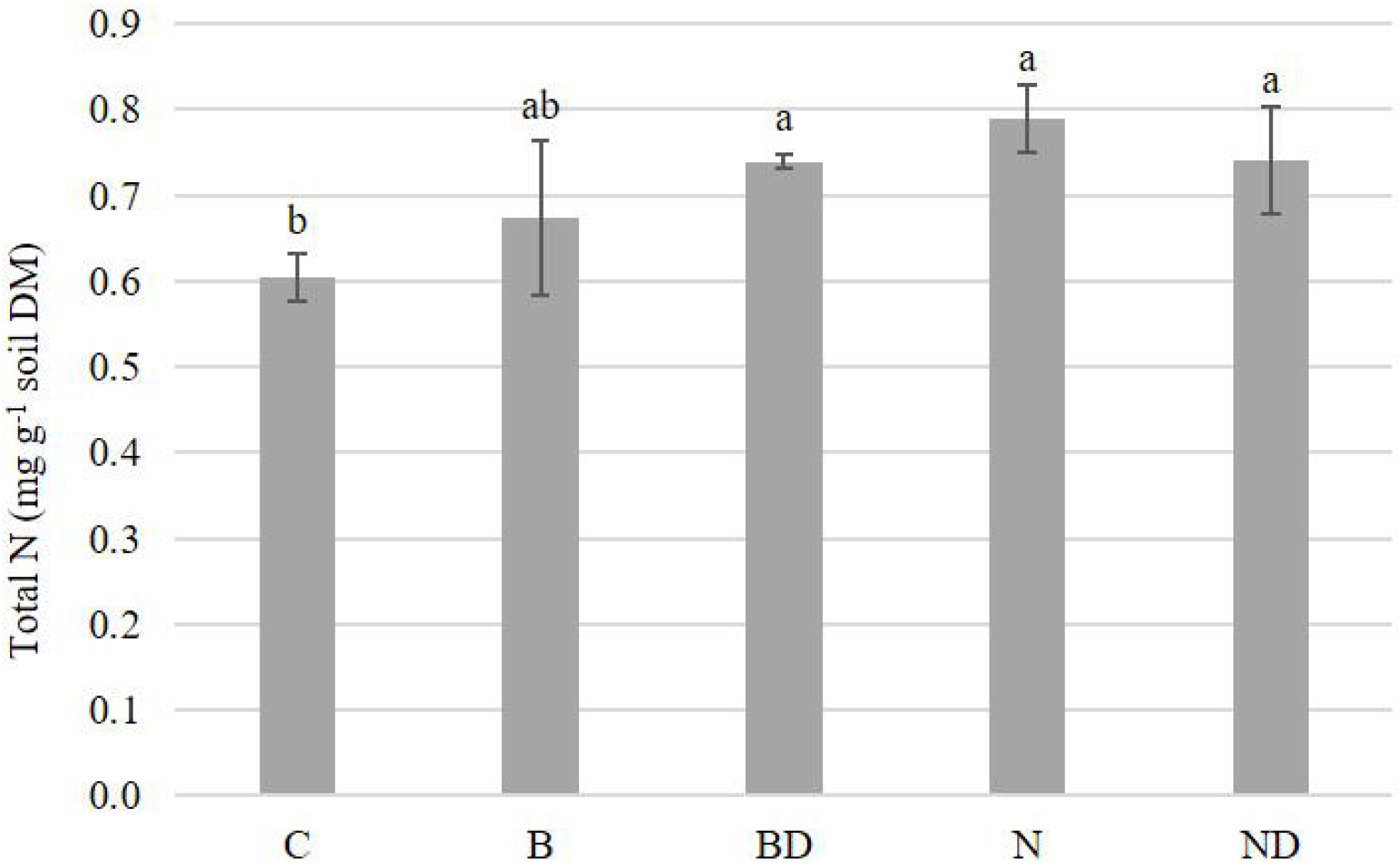

### Phosphorus in biomass/soil

The highest P value has been observed in G1 C plant: 13.8 mg g^-1^ leaves DM **(Figure 6)**. The other lettuce plants state on significantly lower P values comparing to control plant: by 1.6 times in B treatment and by 3.7 times in N treatments and more in the rest of the DAM amended treatments (BD and ND).

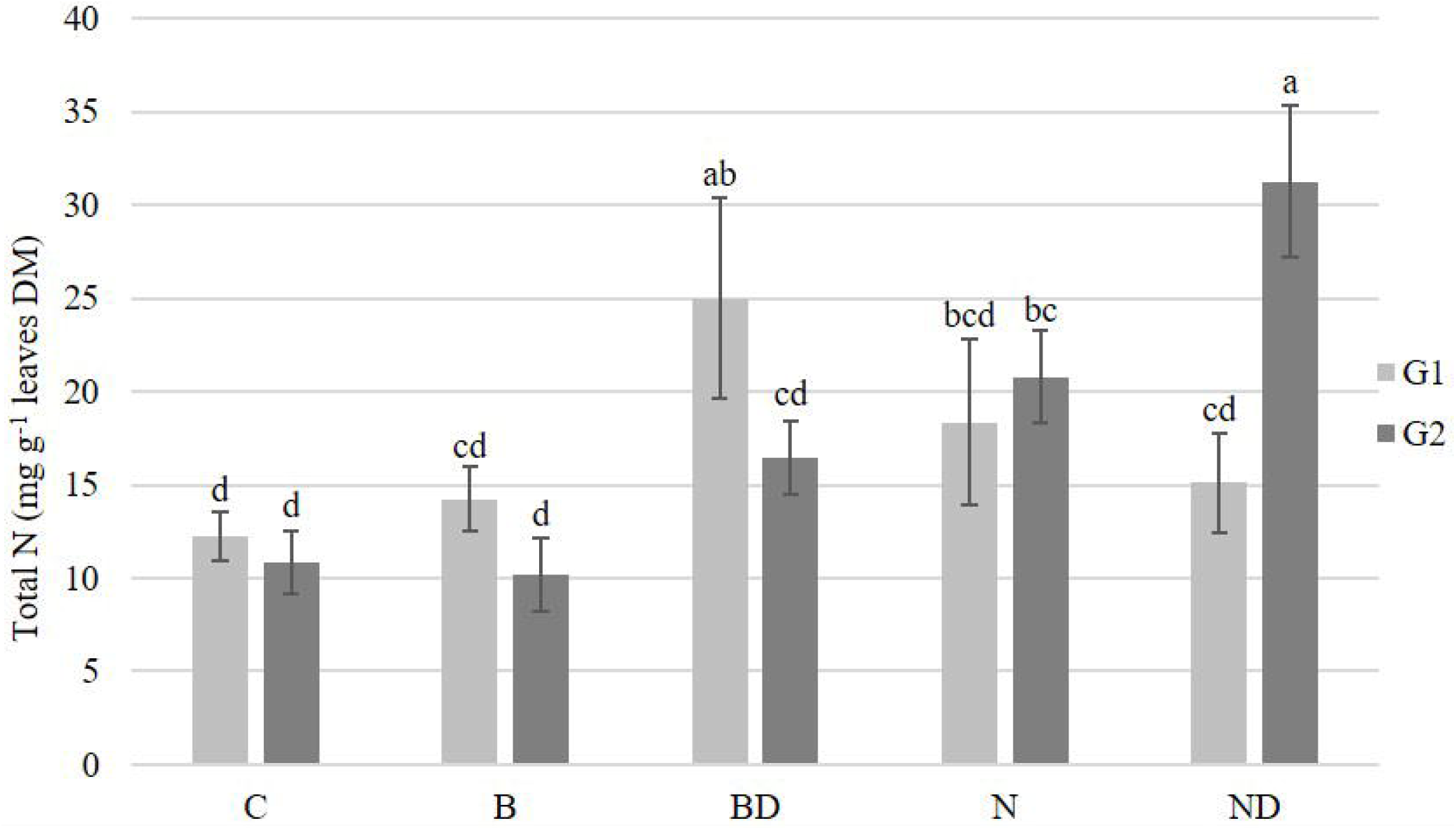

In G1, no statistically significant differences in plant P content have been found between DAM amended treatments (BD and ND) and the Novaferm inoculated plants. In G2, no statistically significant differences have been found within the treated plants including the C plant. P contents varied from 2-2.5 mg g^-1^ leaves DM. Comparing G1 and G2, P concentration decreased in all the treatments of G2, except of the DAM amended ones (BD and ND) having no statistical differences between the growing cycles.

In G2, biomass P content in the control plant dropped by 5.5 times comparing to G1. In B treatment biomass P level has dropped by 3.6 times comparing to G1 and by 1.8 times in N treatment respectively.

No statistically significant differences of soil extractable P have been found between the control soil along with the treated by BCH soils **(Figure 7)**. Values fluctuated from 0.5-0.6 mg g^-1^ soil DM.

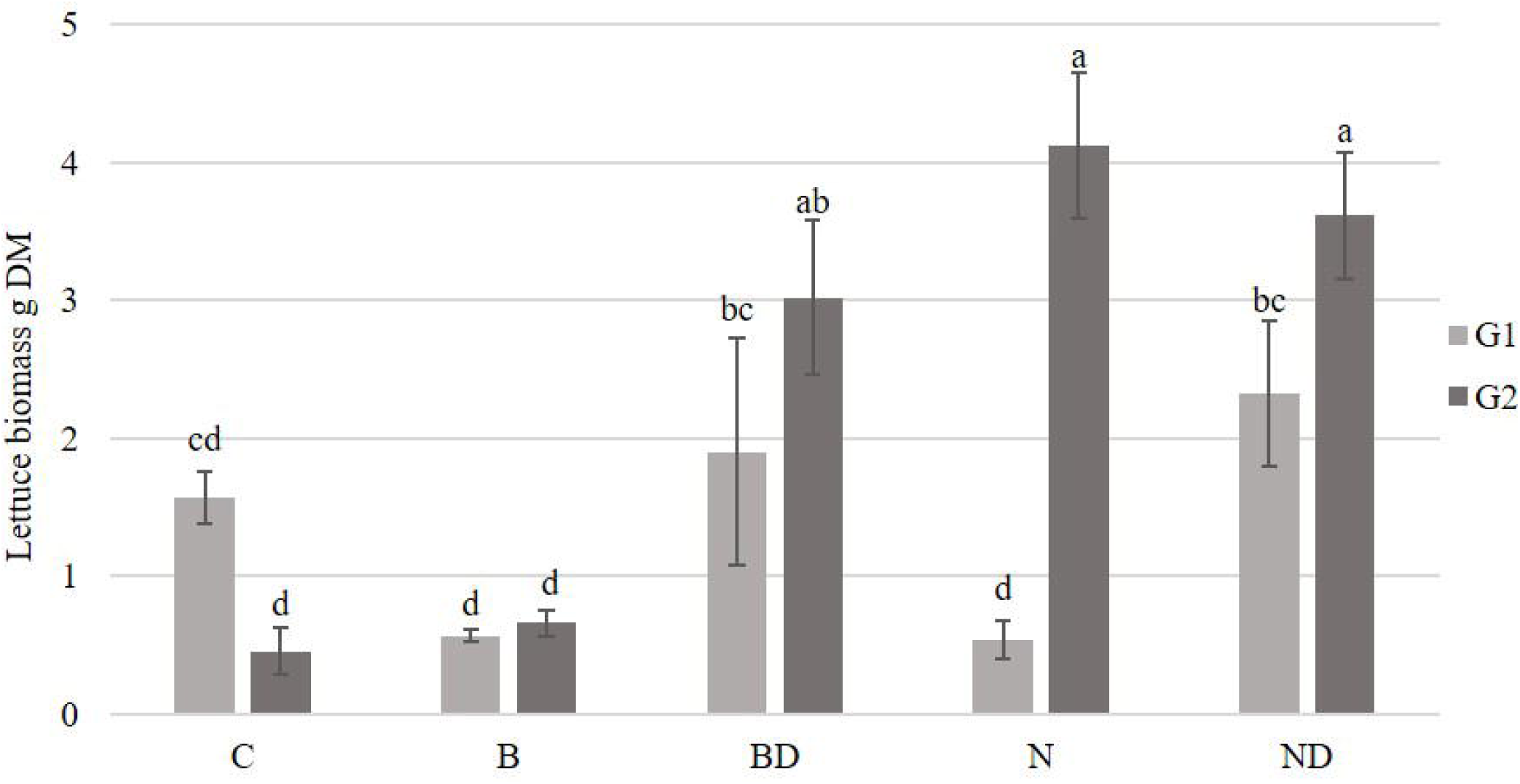

## Discussion

The differences between two plant growing cycles grown in the same BCH amended soil and influenced by the two bacterial inoculums with mineral fertilizer have been investigated throughout the experiment. As hypothesized, the plants from two plant growing cycles distinguished while growing in the same soil, although with the consequently changed soil properties.

### Influence of the BCH mixed with bacterial inoculums

Plant biomass of control plants in G1 and G2 had decreased values comparing to the amended treatments (Figure 1). This might have been caused by the consequent nutrient depletion due to plant growth in G1 and the lack of those nutrients in G2 for the lettuce plants. B plants in G1 and G2 and N plants in G1 had decreased values equal to the control in G2. According to Joseph et al. [31] BCH in high concentrations added into the rhizosphere established contrasting from the typical one environment that would naturally develop there from the typical soil clays, silt, sand and organic matter components, resulting into changed redox potential around the BCH particle. Low biomass N content in control and in Bactofil treatment of both growing cycles has been related to the limitation by the N lack in the soil as well (Figure 2, 3). Initial soil N amount (Table 1) compared to G2 C soil showed a decrease by 62.5% which can be explained by the lettuce nutrient consumption. The results are in the line with the former studies on lettuce growth and sandy soil with compost [32] and with hydroponic solution [33].

The highest plant P content in G1 control plant and the following decreased plant P values in the amended treatments may state on P allocation where in G1 microorganisms in inoculated soils had concurrence for available P. Whereas in the B treated soil no concurrence occurred that consequently resulted into a higher P content in aboveground biomass (Figure 6). Studies of Rodríguez and Fraga [34] on phosphate solubilizing bacteria and P uptake support this hypothesis. From the other side, lower P content in the amended lettuce plants comparing to the higher P ratio of the C plant and B treated plant could have been related to the nutrient dilution within the plant growth resulting into decreased P concentration. Initial soil P content (Table 1) was higher by 64% comparing to the soil P in G2 C soil (Figure 7) might be also explained by the reduced biomass development of C plant in G2 with the decreased P uptake which differ from G1 lettuce plant (Figure 1, 6). The results are in accordance with the studies on lettuce P uptake in a silty clay loam soil [35] and quartz sand with P solutions [36].

Biomass of BCH amended plants with inoculums reduced in G1 (Figure 1) and this might be related to BCH ability to bend nutrients, as demonstrated with burcucumber BCH on lettuce that led to the supressed plant growth [37]. B treated soils showed a decrease in lettuce growth of both G1 and G2 compared to the other treatments. In the studies of Jaiswal et al. [38] on beans and four BCH types (feedstock: eucalyptus wood chips and greenhouse pepper plant wastes) studied “Shifted Rmax-Effect” revealed the effective BCH dose for disease reduction that was lower than that for plant growth promotion, whereas at higher BCH doses no damping-off suppression had been detected, even was promoted in certain cases. From the other hand, Novaferm inoculated soil stimulated lettuce growth in G2 by 87.8% comparing to G1 and to the C plant in G2. This might be explained by the B inoculum type which could need longer time period to fully realize its potential, as its application into the BCH amended soil did not lead to the lettuce development. In addition, biomass values in B treatment of both growing cycles remained unchanged (Figure 1). Our results are in accordance with the studies of Asai et al. [21] where BCH without N fertilizer led to the reduction of rice N uptake and caused decreased grain yield in G1. Decreased N values in the BCH amended treatments with the B inoculum could have been related to N immobilization by BCH and this hypothesis might explain the reduced plant growth in B amended treatments of both growing cycles. Results are in accordance with the experiments on corn stover and hardwood BCH with silt loam soil and corn [39], maize stover BCH and ryegrass with loamy/sandy soil [40]. Otherwise, N content reduction in B treated plants with the reduced biomass might be explained by the B inoculum lower N_2_-fixators activity with its BCH combination comparing to the Novaferm one, with perhaps inefficient microbial population establishment of the added inoculum as suggested in studies of Dempster et al. [41] where jarrah wood BCH supressed microbial development. However, the amount of total soil N (Figure 3) has increased by the application of BCH up to 24% (N treatment), some of which can be explained by the direct input from BCH [42,43][41,42][42,43]. Some portion of N and P is released from the charcoal residues, but these compounds are rather immediately involved into soil-plant nutrient cycle [44]. Soil P allocation without any significant differences among the treatments might be explained by the better phosphate solubilizing bacteria development in the BCH amended soils with inoculums that had the other strategy than releasing P available to plants [45]. Basically, greater P allocation took place within BCH amended soils, as the lettuce development in that case was higher (Figure 1, 7). Another explanation could be related to BCH as an available P adsorbent from the soil as in the studies of Yao et al. [46] on sugar beet biochar removing P from phosphate solution. Consequently, it could have led to the low biomass P values obtained in BCH amended soils of both growing cycles, as the soil mixed with BCH makes P partly unavailable for the plants.

### The effect of the BCH mixed with the bacterial inoculums and DAM fertilizer

BCH combination with inoculums and N fertilizers resulted into a biomass increase in both growing cycles (Figure 1). Overall, practically all the BCH amended treatments showed lettuce biomass increase in G2 (except the upper mentioned B and control). This trend confirms our hypothesis of BCH persistence in soil and consequent pores inoculation in course of time that drives to plant better growth and therefore biomass increase as suggested previously [47,48]. DAM fertilizer combined with inoculums also initiated total N content increase in biomass of BD treatment comparing to the C (Figure 2). This can be explained by the application of N additives into the soil and thus N availability that correlates with increased biomass growth [49]. Plant N content of ND treatment was not significantly different in G1 compared to C, that might be due to microbial N immobilization as in the studied N microsites suggested by Schimel and Bennett [50]. In the case of ND treatment in G1 the tendency of a decreased plant N amounts up to 48% comparing to G2 could have stated on its use primarily by soil bacteria in G1, as suggested by Kaye and Hart [51] considering N competition within plants and soil microorganisms. Soil N content remained unchanged between BCH amended treatments (Figure 3), but was significantly higher than the C soil (treatments BD, N and ND). BCH may also limit soil N availability in N deficient soils due to the high C/N ratio and temporarily reduce crop productivity [52].

Hence, significantly raised Corg values have been found in all the BCH treated soils with the highest value in BD treated soil by 80% (Figure 5). Obtained results where the Corg doubled in the BCH amended soils are confirmed as well by the studies of Lehmann et al. [53] involving Ferrasol and using cowpea as a test plant. Other studies confirm, that for example maize BCH addition in augmented amounts led to great soil organic carbon (SOC) rise and total N as well [54]. Moreover, BCH addition associated with the increased SOC contents might enhance the nutrient retention capacity of the soil due to the higher CEC with organo-mineral complexes forming [55]. From the other side, BCH application in high rates like 16 t ha^-1^ caused N limitation, even with N fertilizer addition, and thus low grain yields with the reduction of rice plant N uptake [21]. In the other studies, maize BCH and urea fertilizer lead to the short-term reductions in soil mineral N availability as a result of probable BCH negative effect on soil quality and fertility characteristics [54]. It might be also caused by the significant absorption of N up to 22.1% with the ash/charcoal woody residues according to the studies of Dünisch et al. [56], where actually the same has been observed in the case of available P compounds (up to 11.7%). Biomass and soil P content remained without any significant differences between the G2 treatments (Figure 6, 7). The explanation can be found in the blocked soil P due to the increased pH that is caused by BCH amendment [57]. BCH absorbs P and also increases P fertilizer retention in soils, though its acting intensity broadly depends on BCH feedstock type [58] or can also adsorb phosphate efficiently from solutions being a potential P source [59]. The other investigation, that studied the influence of BCH and N fertilizers on different soil types from several locations in northern Laos, found yield rise in soils with low P availability and also enhanced plant reaction to additional fertilizers with BCH additions [21].

## Conclusion

The conducted research proves the importance of BCH enrichment during persistence in soil and its consequent positive influence on plant growth after certain period of time. The resulted nutrient depletion by plant can be compensated by the BCH bacterial inoculation and its mixing with the mineral fertilizer. Making the comparison between two bacterial additives mixed with BCH and their persistence in soil it has been concluded, that growth promoting effect of inoculums was supported by N fertilization. Regarding the influence on the total N content in plant, Novaferm bacteria diversity promotes better N availability and assimilation possibility, and consequently the plant growth. Thus, Novaferm inoculum tends to increase N bonding in G2 via bacterial N_2_-fixation in greater extent than Bactofil. BCH most probably decreases N availability in soil. Higher C and P content in all biochar amended soils should be attributed to C and nutrient input in added BCH. Nevertheless, in both inoculums Bactofil and Novaferm are phosphate solubilizing bacteria not efficient enough. This investigation needs deeper study aiming to analyse BCH aging in soil for a longer period and its functioning in soil-plant nutrient cycling.

## Supporting information

Acknowledgements

Supplemental list of figures and tables

## Acknowledgements

This paper was supported by the project ATCZ42 INTEKO: “Innovation technologies in composting, its application and soil protection”; by the Internal Grant Agency IGA FA MENDELU (Faculty of Agriscience, Mendel University in Brno) No. TP 3/2015 with the support of the Specific University Research Grant (Ministry of Education, Youth and Sports, Czech Republic); by the Erasmus+ mobility program realizing the internship at ISA, Lille (France).

## References

[1] Tilman D, Cassman KG, Matson PA, et al. Agricultural sustainability and intensive production practices. Nature. 2002;418:671–677.

[2] Lal R. Anthropogenic Influences on World Soils and Implications to Global Food Security. Adv. Agron. 2007;93:69–93.

[3] Osman KT. Soil degradation, conservation and remediation. Soil Degrad. Conserv. Remediat. 2014.

[4] Prazan J, Dumbrovsky M. Soil conservation policies: Conditions for their effectiveness in the Czech Republic. L. Degrad. Dev. 2011;22:124–133.

[5] Lal R. Soil Carbon Sequestration Impacts on Global Climate Change and Food Security. Am. Assoc. Adv. Sci. 2004;304:1623–1627.

[6] Richter DDB. Humanity’s transformation of earth’s soil: Pedology’s new frontier. Soil Sci. 2007;172:957–967.

[7] Lehmann J, Gaunt J, Rondon M. Bio-char sequestration in terrestrial ecosystems - A review [Internet]. Mitig. Adapt. Strateg. Glob. Chang. Kluwer Academic Publishers-Plenum Publishers; 2006 [cited 2017 Jun 2]. p. 403–427. Available from: http://link.springer.com/10.1007/s11027-005-9006-5.

[8] Major J, Rondon M, Molina D, et al. Maize yield and nutrition during 4 years after biochar application to a Colombian savanna oxisol. Plant Soil [Internet]. 2010 [cited 2018 Feb 8];333:117–128. Available from: https://link.springer.com/content/pdf/10.1007%2Fs11104-010-0327-0.pdf.

[9] Spokas KA, Cantrell KB, Novak JM, et al. Biochar: A Synthesis of Its Agronomic Impact beyond Carbon Sequestration. J. Environ. Qual. [Internet]. 2012;41:973. Available from: https://www.agronomy.org/publications/jeq/abstracts/41/4/973.

[10] Wiedner K, Schneewei J, Dippold MA, et al. Anthropogenic Dark Earth in Northern Germany - The Nordic Analogue to terra preta de Índio in Amazonia. Catena. 2014;132:114–125.

[11] Rondon MA, Lehmann J, Ramírez J, et al. Biological nitrogen fixation by common beans (Phaseolus vulgaris L.) increases with bio-char additions. Biol. Fertil. Soils [Internet]. 2007 [cited 2018 Mar 7];43:699–708. Available from: http://link.springer.com/10.1007/s00374-006-0152-z.

[12] Warnock DD, Lehmann J, Kuyper TW, et al. Mycorrhizal responses to biochar in soil - Concepts and mechanisms [Internet]. Plant Soil. 2007 [cited 2017 Jun 2]. p. 9–20. Available from: http://download.springer.com/static/pdf/360/art%253A10.1007%252Fs11104-007-9391-5.pdf?originUrl=http%3A%2F%2Flink.springer.com%2Farticle%2F10.1007%2Fs11104-007-9391-5&token2=exp=1496421697∼acl=%2Fstatic%2Fpdf%2F360%2Fart%25253A10.1007%25252Fs11104-007-939.

[13] Martínez-Viveros O, Jorquera M., Crowley D., et al. Mechanisms and Practical Considerations Involved in Plant Growth Promotion By Rhizobacteria. J. soil Sci. plant Nutr. [Internet]. 2010 [cited 2018 Jun 3];10:293–319. Available from: http://www.scielo.cl/scielo.php?script=sci_arttext&pid=S0718-95162010000100006&lng=en&nrm=iso&tlng=en.

[14] Warnock DD, Mummey DL, McBride B, et al. Influences of non-herbaceous biochar on arbuscular mycorrhizal fungal abundances in roots and soils: Results from growth-chamber and field experiments. Appl. Soil Ecol. [Internet]. 2010 [cited 2018 Jun 3];46:450–456. Available from: https://www.sciencedirect.com/science/article/pii/S0929139310001654.

[15] Zimmerman AR, Gao B, Ahn MY. Positive and negative carbon mineralization priming effects among a variety of biochar-amended soils. Soil Biol. Biochem. [Internet]. 2011;43:1169–1179. Available from: http://dx.doi.org/10.1016/j.soilbio.2011.02.005.

[16] Deenik JL, McClellan T, Uehara G, et al. Charcoal Volatile Matter Content Influences Plant Growth and Soil Nitrogen Transformations. Soil Sci. Soc. Am. J. [Internet]. 2010;74:1259. Available from: https://www.soils.org/publications/sssaj/abstracts/74/4/1259.

[17] Gundale MJ, DeLuca TH. Charcoal effects on soil solution chemistry and growth of Koeleria macrantha in the ponderosa pine/Douglas-fir ecosystem. Biol. Fertil. Soils [Internet]. 2007 [cited 2018 May 31];43:303–311. Available from: http://link.springer.com/10.1007/s00374-006-0106-5.

[18] Liang B, Lehmann J, Solomon D, et al. Black Carbon Increases Cation Exchange Capacity in Soils. Soil Sci. Soc. Am. J. [Internet]. 2006;70:1719. Available from: https://www.soils.org/publications/sssaj/abstracts/70/5/1719.

[19] Yao FX, Arbestain MC, Virgel S, et al. Simulated geochemical weathering of a mineral ash-rich biochar in a modified Soxhlet reactor. Chemosphere [Internet]. 2010 [cited 2018 Apr 22];80:724–732. Available from: https://www.sciencedirect.com/science/article/pii/S0045653510006028.

[20] Verheijen F, Jeffery S, Bastos a C, et al. Biochar application to soils: a critical review of effects on soil properties, processes and functions. JRC Sci. Tech. Rep. 2010.

[21] Asai H, Samson BK, Stephan HM, et al. Biochar amendment techniques for upland rice production in Northern Laos. 1. Soil physical properties, leaf SPAD and grain yield. F. Crop. Res. [Internet]. 2009 [cited 2018 May 27];111:81–84. Available from: https://www.sciencedirect.com/science/article/pii/S0378429008002141.

[22] Chan KY, Van Zwieten L, Meszaros I, et al. Agronomic values of green waste biochar as a soil amendment. Aust. J. Soil Res. 2007;45:629–634.

[23] Steiner C, Teixeira WG, Lehmann J, et al. Long term effects of manure, charcoal and mineral fertilization on crop production and fertility on a highly weathered Central Amazonian upland soil. Plant Soil [Internet]. 2007 [cited 2018 Jan 22];291:275–290. Available from: https://link.springer.com/content/pdf/10.1007%2Fs11104-007-9193-9.pdf.

[24] Chia CH, Singh BP, Joseph S, et al. Characterization of an enriched biochar. J. Anal. Appl. Pyrolysis [Internet]. 2014 [cited 2018 May 6];108:26–34. Available from: https://www.sciencedirect.com/science/article/pii/S0165237014001272.

[25] Fazal A, Bano A. Role of Plant Growth-Promoting Rhizobacteria (PGPR), Biochar, and Chemical Fertilizer under Salinity Stress. Commun. Soil Sci. Plant Anal. [Internet]. 2016;47:1985–1993. Available from: http://dx.doi.org/10.1080/00103624.2016.1216562.

[26] Conversa G, Bonasia A, Lazzizera C, et al. Influence of biochar, mycorrhizal inoculation, and fertilizer rate on growth and flowering of Pelargonium (Pelargonium zonale L.) plants. Front. Plant Sci. [Internet]. 2015 [cited 2019 Apr 16];6:429. Available from: http://journal.frontiersin.org/Article/10.3389/fpls.2015.00429/abstract.

[27] Pokorný E, Šarapatka B, Hejátková K. Hodnocení kvality půdy v ekologicky hospodařícím podniku. ZERA, Náměšt and Oslavou; 2007.

[28] Hammes K, Smernik RJ, Skjemstad JO, et al. Synthesis and characterisation of laboratory-charred grass straw (Oryza sativa) and chestnut wood (Castanea sativa) as reference materials for black carbon quantification. Org. Geochem. [Internet]. 2006 [cited 2019 Feb 17];37:1629–1633. Available from: https://www.sciencedirect.com/science/article/pii/S0146638006001653.

[29] Saha UK, Sonon L, Kissel DE. Comparison of Conductimetric and Colorimetric Methods with Distillation–Titration Method of Analyzing Ammonium Nitrogen in Total Kjeldahl Digests. Commun. Soil Sci. Plant Anal. 2012;43:2323–2341.

[30] Joret F, Hébert J. Contribution à la détermination du besoin des sols en acide phosphorique. Ann. Agron. 1955;233–299.

[31] Joseph S, Husson O, Graber ER, et al. The electrochemical properties of biochars and how they affect soil redox properties and processes [Internet]. Agronomy. Multidisciplinary Digital Publishing Institute; 2015 [cited 2019 Sep 6]. p. 322–340. Available from: http://www.mdpi.com/2073-4395/5/3/322.

[32] Brito LM. Lettuce (Lactuca sativa L.) and Cabbage (Brassica oleracea L. var. capitata L.) Growth in Soil Mixed with Municipal Solid Waste Compost and Paper Mill Sludge Composted with Bark. Acta Hortic. 2001;563:131–137.

[33] Domingues DS, Takahashi HW, Camara CAP, et al. Automated system developed to control pH and concentration of nutrient solution evaluated in hydroponic lettuce production. Comput. Electron. Agric. [Internet]. 2012 [cited 2019 Jul 7];84:53–61. Available from: https://www.sciencedirect.com/science/article/pii/S0168169912000361.

[34] Rodríguez H, Fraga R. Phosphate solubilizing bacteria and their role in plant growth promotion. Biotechnol. Adv. [Internet]. 1999 [cited 2019 Jul 7];17:319–339. Available from: https://www.sciencedirect.com/science/article/pii/S0734975099000142.

[35] Chabot R, Antoun H, Cescas MP. Growth promotion of maize and lettuce by phosphate-solubilizing Rhizobium leguminosarum biovar. phaseoli. Plant Soil. 1996;184:311–321.

[36] Xu G, Levkovitch I, Soriano S, et al. Integrated effect of irrigation frequency and phosphorus level on lettuce: P uptake, root growth and yield. Plant Soil [Internet]. 2004 [cited 2019 Jul 14];263:297–309. Available from: http://link.springer.com/10.1023/B:PLSO.0000047743.19391.42.

[37] Rajapaksha AU, Vithanage M, Lim JE, et al. Invasive plant-derived biochar inhibits sulfamethazine uptake by lettuce in soil. Chemosphere [Internet]. 2014 [cited 2017 Aug 17];111:500–504. Available from: http://ac.els-cdn.com/S0045653514005232/1-s2.0-S0045653514005232-main.pdf?_tid=dfe8e1a6-8374-11e7-8fa2-00000aab0f26&acdnat=1502992547_5f9db38698edb56d45315fa443f186cc.

[38] Jaiswal AK, Frenkel O, Elad Y, et al. Non-monotonic influence of biochar dose on bean seedling growth and susceptibility to Rhizoctonia solani: the “Shifted Rmax-Effect.” Plant Soil [Internet]. 2015 [cited 2019 Sep 6];395:125–140. Available from: http://link.springer.com/10.1007/s11104-014-2331-2.

[39] Fidel RB, Laird DA, Parkin TB. Impact of six lignocellulosic biochars on C and N dynamics of two contrasting soils. GCB Bioenergy [Internet]. 2017 [cited 2019 Mar 7];9:1279–1291. Available from: http://doi.wiley.com/10.1111/gcbb.12414.

[40] Liu Z, He T, Cao T, et al. Effects of biochar application on nitrogen leaching, ammonia volatilization and nitrogen use efficiency in two distinct soils. J. soil Sci. plant Nutr. [Internet]. 2017 [cited 2019 Mar 1];17:0–0. Available from: http://www.scielo.cl/scielo.php?script=sci_arttext&pid=S0718-95162017005000037&lng=en&nrm=iso&tlng=en.

[41] Dempster DN, Gleeson DB, Solaiman ZM, et al. Decreased soil microbial biomass and nitrogen mineralisation with Eucalyptus biochar addition to a coarse textured soil. Plant Soil. 2012;354:311–324.

[42] Nguyen TTN, Xu CY, Tahmasbian I, et al. Effects of biochar on soil available inorganic nitrogen: A review and meta-analysis [Internet]. Geoderma. Elsevier; 2017 [cited 2018 May 5]. p. 79–96. Available from: https://www.sciencedirect.com/science/article/pii/S0016706116307388?via%3Dihub.

[43] Zhang H, Voroney RP, Price GW. Effects of temperature and processing conditions on biochar chemical properties and their influence on soil C and N transformations. Soil Biol. Biochem. [Internet]. 2015;83:19–28. Available from: http://dx.doi.org/10.1016/j.soilbio.2015.01.006.

[44] Gul S, Whalen JK. Biochemical cycling of nitrogen and phosphorus in biochar-amended soils. Soil Biol. Biochem. [Internet]. 2016;103:1–15. Available from: http://dx.doi.org/10.1016/j.soilbio.2016.08.001.

[45] Jilani G, Saqlan Naqvi SM, Rasheed MH, et al. Phosphorus Solubilizing Bacteria: Occurrence, Mechanisms and their Role in Crop Production Breeding for oil quality View project exploration of in planta Agrobacterium mediated transformation in lentil (Lens culinaris Medik) View project. 2009; Available from: https://www.researchgate.net/publication/307108426.

[46] Yao Y, Gao B, Inyang M, et al. Bioresource Technology Biochar derived from anaerobically digested sugar beet tailings[: Characterization and phosphate removal potential. Bioresour. Technol. [Internet]. 2011;102:6273–6278. Available from: http://dx.doi.org/10.1016/j.biortech.2011.03.006.

[47] Jones DL, Rousk J, Edwards-Jones G, et al. Biochar-mediated changes in soil quality and plant growth in a three year field trial. Soil Biol. Biochem. [Internet]. 2012 [cited 2018 Sep 26];45:113–124. Available from: https://ac.els-cdn.com/S0038071711003865/1-s2.0-S0038071711003865-main.pdf?_tid=e0b8475c-efec-4dd2-8d6b-ac406d6b4b85&acdnat=1537999107_dda086655608fd8e2e0384e7a5a47cfd.

[48] Biederman LA, Stanley Harpole W. Biochar and its effects on plant productivity and nutrient cycling: A meta-analysis. GCB Bioenergy. 2013;5:202–214.

[49] Liu X, Xu J, Yu L, et al. Combined application of biochar and nitrogen fertilizer benefits nitrogen retention in the rhizosphere of soybean by increasing microbial biomass but not altering microbial community structure. Sci. Total Environ. [Internet]. 2018 [cited 2019 Mar 1];640–641:1221–1230. Available from: https://www.sciencedirect.com/science/article/pii/S0048969718320801?via%3Dihub.

[50] Schimel JP, Bennett J. Nitrogen mineralization: challenges of a changing paradigm. Ecology. 2004;85(3):591–602.

[51] Kaye JP, Hart SC. Competition for nitrogen between plants and soil microorganisms [Internet]. Trends Ecol. Evol. Elsevier Current Trends; 1997 [cited 2019 Jul 8]. p. 139–143. Available from: https://www.sciencedirect.com/science/article/pii/S016953479701001X.

[52] Lehmann J, Kern D, German L, et al. Soil Fertility and Production Potential. Amaz. Dark Earths [Internet]. Dordrecht: Kluwer Academic Publishers; 2003 [cited 2018 Jan 20]. p. 105–124. Available from: http://link.springer.com/10.1007/1-4020-2597-1_6.

[53] Lehmann J, Pereira da Silva J, Steiner C, et al. Nutrient availability and leaching in an archaeological Anthrosol and a\rFerralsol of the Central Amazon basin: fertilizer, manure and charcoal\ramendments. Plant Soil [Internet]. 2003 [cited 2018 May 27];249:343–357. Available from: https://link.springer.com/content/pdf/10.1023%2FA%3A1022833116184.pdf.

[54] Wang X, Song D, Liang G, et al. Maize biochar addition rate influences soil enzyme activity and microbial community composition in a fluvo-aquic soil. Appl. Soil Ecol. [Internet]. 2015;96:265–272. Available from: http://dx.doi.org/10.1016/j.apsoil.2015.08.018.

[55] Glaser B, Lehmann J, Zech W. Ameliorating physical and chemical properties of highly weathered soils in the tropics with charcoal - A review. Biol. Fertil. Soils. 2002;35:219–230.

[56] Dünisch O, Lima VC, Seehann G, et al. Retention properties of wood residues and their potential for soil amelioration. Wood Sci. Technol. [Internet]. 2007 [cited 2018 May 7];41:169–189. Available from: http://link.springer.com/10.1007/s00226-006-0098-1.

[57] Takaya CA, Fletcher LA, Singh S, et al. Phosphate and ammonium sorption capacity of biochar and hydrochar from different wastes. Chemosphere [Internet]. 2016 [cited 2018 Feb 7];145:518–527. Available from: https://ac.els-cdn.com/S0045653515303866/1-s2.0-S0045653515303866-main.pdf?_tid=be1cb7ee-0c1f-11e8-82f1-00000aacb362&acdnat=1518019291_60a66c2f0e9bf79a0fc816e93168383f.

[58] Zhang H, Chen C, Gray EM, et al. Roles of biochar in improving phosphorus availability in soils: A phosphate adsorbent and a source of available phosphorus. Geoderma [Internet]. 2016 [cited 2018 May 28];276:1–6. Available from: http://linkinghub.elsevier.com/retrieve/pii/S0016706116301793.

[59] Liang X, Jin Y, He M, et al. Phosphorus speciation and release kinetics of swine manure biochar under various pyrolysis temperatures. Environ. Sci. Pollut. Res. [Internet]. 2017 [cited 2018 Feb 13];1–9. Available from: https://link.springer.com/content/pdf/10.1007%2Fs11356-017-0640-8.pdf.

