## Supplemental list of figures and tables for "Plant-soil nitrogen, carbon and phosphorus content after the addition of biochar, bacterial inoculums and N fertilizer"

**Table 1.** Banín soil content characteristics

**Table 2.** Physicochemical characteristics of studied biochar

**Table 3.** Characteristics of all the applied treatments

**Figure 1.** Aboveground biomass production (g DM) within two plant growing cycles – the first (G1) and the second (G2) in pots with treatments (C, B, BD, N and ND). Values are presented as means ± SD. Different letters refer to significant differences between plants (Tukey HSD test, n = 5, p ≤ 0.05).

**Figure 2.** Nitrogen content (mg g^-1^ leaves DM) in aboveground lettuce biomass of the first (G1) and the second (G2) growing cycles in five different soil treatments (C, B, BD, N and ND). Values are presented as means ± SD. Different letters refer to significant differences between plants (Tukey HSD test, n = 4, p ≤ 0.05).

**Figure 3.** Total soil N content (mg g^-1^ soil DM) after G2 plants harvesting in five different soil treatments (C, B, BD, N and ND). Values are presented as means ± SD. Different letters refer to significant differences between plants (Tukey HSD test, n = 4, p ≤ 0.05).

**Figure 4.** Corg content (mg g^-1^ leaves DM) in aboveground lettuce biomass of the first (G1) and the second (G2) growing cycles in five different soil treatments (C, B, BD, N and ND). Values are presented as means ± SD. Different letters refer to significant differences between plants (Tukey HSD test, n = 4, p ≤ 0.05).

**Figure 5.** Soil Corg content (mg g^-1^ soil DM) after G2 plants harvesting in five different soil treatments (C, B, BD, N and ND). Values are presented as means ± SD. Different letters refer to significant differences between plants (Tukey HSD test, n = 4, p ≤ 0.05).

**Figure 6.** P content (mg g^-1^ leaves DM) in aboveground lettuce biomass of the first (G1) and the second (G2) growing cycles in five different soil treatments (C, B, BD, N and ND). Values are presented as means ± SD. Different letters refer to significant differences between plants (Tukey HSD test, n = 4, p ≤ 0.05).

**Figure 7.** Soil P content (mg g^-1^ soil DM) of second plant growing cycle (G2) in five different soil treatments (C, B, BD, N and ND). Values are presented as means ± SD. Different letters refer to significant differences between plants (Tukey HSD test, n = 4, p ≤ 0.05).
